# A single cell atlas of the mouse seminal vesicle

**DOI:** 10.1101/2024.04.08.588538

**Authors:** Fengyun Sun, Kathleen Desevin, Yu Fu, Shanmathi Parameswaran, Jemma Mayall, Vera Rinaldi, Nils Krietenstein, Artur Manukyan, Qiangzong Yin, Carolina Galan, Chih-Hsiang Yang, Anastasia V. Shindyapina, Vadim N. Gladyshev, Manuel Garber, John E. Schjenken, Oliver J. Rando

## Abstract

During mammalian reproduction, sperm are delivered to the female reproductive tract bathed in a complex medium known as seminal fluid, which plays key roles in signaling to the female reproductive tract and in nourishing sperm for their onwards journey. Along with minor contributions from the prostate and the epididymis, the majority of seminal fluid is produced by a somewhat understudied organ known as the seminal vesicle. Here, we report the first single-cell RNA-seq atlas of the mouse seminal vesicle, generated using tissues obtained from 23 mice of varying ages, exposed to a range of dietary challenges. We define the transcriptome of the secretory cells in this tissue, identifying a relatively homogeneous population of the epithelial cells which are responsible for producing the majority of seminal fluid. We also define the immune cell populations – including large populations of macrophages, dendritic cells, T cells, and NKT cells – which have the potential to play roles in producing various immune mediators present in seminal plasma. Together, our data provide a resource for understanding the composition of an understudied reproductive tissue with potential implications for paternal control of offspring development and metabolism.

## INTRODUCTION

Although the core event in reproduction of sexually-reproducing species is the merging of two haploid genomes to re-generate a diploid genome, a variety of other processes support efficient reproduction, for instance provisioning gametes to support them during the at times lengthy process of finding one another. In internally-fertilizing species, sperm are delivered to the female reproductive tract carried in a complex solution known as seminal fluid, which has well-known roles in gamete nutrition but is increasingly understood to also carry out an ever-expanding suite of signaling functions ^1–5^. In mammals, immune modulators (transforming growth factor beta family members, E type prostaglandins, cytokines, etc.) present in seminal fluid help prepare the endometrium to be receptive to implantation ^2,3,5,6^. Seminal fluid has also been implicated in modulating offspring phenotypes: in a milestone study, Bromfield *et al* transferred embryos to females mated with vasectomized males with or without seminal vesicles, and characterized offspring phenotypes ^7^. Male offspring born in the absence of seminal vesicle secretions exhibited dramatic metabolic defects, including increased adiposity and decreased glucose tolerance.

Moreover, the contents of seminal fluid can be modulated by paternal lifestyle ^1,8–13^; although several studies have shown that some effects of paternal diet or stress on offspring can be transmitted by sperm, Watkins *et al* reported that *both* sperm and seminal fluid from low protein-fed males independently modulate offspring cardiovascular health ^1,10^, while Lassi *et al* reported that paternal effects of circadian disruption were likely mediated entirely by seminal fluid ^11^. Other studies have reported dietary and other effects on seminal fluid cytokine composition ^8,9^ – accompanied by changes to maternal reproductive tract gene regulation ^8^ – albeit without linking these changes to offspring phenotypes.

The majority (∼80%, in mouse) of seminal plasma is contributed by a male accessory reproductive organ known as the seminal vesicle, with some (∼15%) additional contribution from the prostate. Unlike the prostate, which has been intensively characterized thanks to the prevalence of prostate cancer in humans, the seminal vesicle is relatively understudied. Histologically, the seminal vesicle is comprised of a single layer of secretory epithelial cells organized in a pseudostratified columnar epithelium surrounding a lumen filled with seminal plasma. The basal aspect of this epithelial sheet is surrounded by a layer of flat basal cells similar histologically to those found in other tissues such as the epididymis and kidney. Supporting stromal cells include fibroblasts and smooth muscle – whose contraction drives seminal fluid into the ejaculatory duct – as well as endothelial cells and infiltrating immune cells.

In the past few years, systematic efforts have explored gene regulation ^13,14^ and proteomics ^12,15^ of the seminal vesicle tissue, along with proteomics of seminal vesicle fluid ^9,16–19^. These studies have highlighted important signaling pathways that likely influence seminal vesicle secretory function. However, given the complex cellular profile of seminal vesicle tissue, the specific cell types that drive these secretions remain to be identified. Over the past decade, advances in low input genomics methods such as single cell RNA-seq have enabled rapid and systematic characterization of cell composition and cell type-specific gene regulation in complex tissues ^20,21^. To our knowledge the seminal vesicle has not yet been characterized by single cell RNA-seq, making it one of the last – perhaps the last – major mammalian organ system lacking a single cell atlas. We therefore set out here to characterize the cell composition of this tissue by single cell RNA-seq, both to establish gene expression in the various cell types of this key reproductive tissue, and to potentially to identify previously unappreciated subpopulations of various cell types.

## RESULTS

### Tissue collection and single cell RNA-seq

We set out to generate a single cell RNA-seq atlas of the murine seminal vesicle to define cell types comprising this tissue and to characterize the gene expression program for the various cell populations present. A typical tissue preparation is shown in **Figure S1**. As seminal fluid composition can be influenced by diets and other stressors, we collected tissues from animals across a range of ages (from ∼12 weeks to ∼28 months of age) and subject to various dietary challenges from caloric restriction to high fat diet exposure (**Methods**). Altogether, we generated data for a total of 23 samples (**Table S1**), using the 10X Genomics microfluidic platform for cell barcoding. After quality control measures (removing empty drops and low quality cells) we captured a total of 47,804 cells with an average of 6053 (median: 2402) unique molecular identifiers (UMIs) per cell. As the seminal vesicle is one of the last major tissues in mammals which has not been subject to single cell RNA-seq, we focus primarily on detailing the major cell types revealed in the complete dataset; dietary and age effects on the seminal vesicle will be explored in followup efforts, as we do not have sufficient cells for any individual exposure in this dataset to robustly identify diet or age-specific effects on this tissue.

### Overview of cell composition of the mouse seminal vesicle

To visualize the cell types that comprise the seminal vesicle we clustered all 47,804 cells and visualized cell populations using the Uniform Manifold Approximation and Projection (UMAP) visualization. We find seven large clusters of cells (**Figure 1A**), which we annotate based on marker genes (**Figures 1B-C**) and which correspond to secretory epithelial cells (with a primary population defined by *Svs1*, *Svs5*, etc., along with one unanticipated population of prostate-like cells; see below), basal cells (*Cldn4*, *Krt8*), stromal cells including fibroblasts (*Col6a5*, *Col1a1*, *Gsn*), smooth muscle (*Tpm2*, *Myh11*), and endothelial cells (*Emcn*, *Pecam1*). Additionally, we observe three groups of immune cells including a cluster comprised of macrophages (*Ctsb*, *Ccl3*), a cluster of monocytes/dendritic cells (*Cd74*, *Xcr1*), and a cluster of T (*Cd3g*, *Cd28*, *Lck*) and presumptive NKT cells (*Cd3g*, *Gzma*). Several smaller clusters are also identified, corresponding to lymphatic endothelial cells, pericytes, and Schwann cells. Altogether, these cell types include all the major cell types found in histological studies ^22–26^. Below, we isolate each of the major cell types for reclustering to explore possible specialization within each cell population.

**Figure 1.**
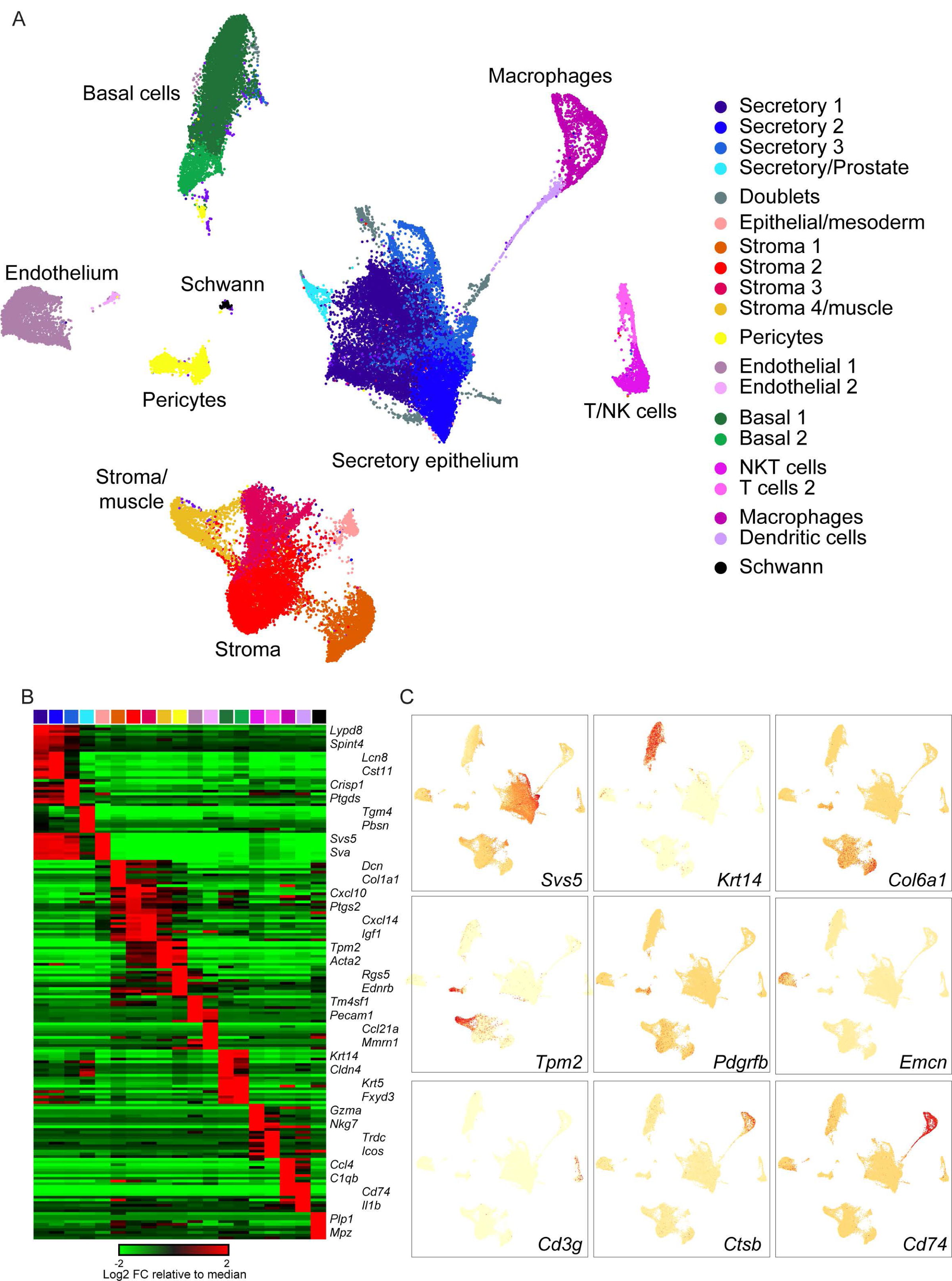
Major cell populations comprising the murine seminal vesicle. A) Overview of full dataset. UMAP visualization of all cells in the dataset, annotated according to inferred cell type. B) Heatmap showing the twenty most enriched genes for each of the 21 clusters in the full dataset. C) UMAPs colored according to expression of marker genes for major cell types in the seminal vesicle.

### Support cells: fibroblasts, smooth muscle, endothelium, and basal epithelial cells

We begin by briefly discussing the various support cells which are common to many or most multicellular tissues and are identified in our seminal vesicle dataset (**Figure 1**). These include three large clusters, each comprised of several subpopulations: stromal cells (fibroblasts and muscle), basal epithelial cells, and endothelial cells. Other support cell types included two smaller clusters, representing pericytes – contractile cells that envelop capillaries in many tissues – and Schwann cells, a type of glial cell which myelinates peripheral neuronal processes. Overall, we find little evidence for meaningful heterogeneity among the populations of basal cells, endothelium, pericytes, or Schwann cells; and as such, we do not further discuss them here.

In contrast, there was clear evidence for heterogeneity in the large group of stromal/fibroblast/muscle cells, motivating us to recluster these cell populations to explore any diversity in cell populations that might be obscured in the whole tissue visualization (**Figure S2**). Although we did not identify well-separated clusters, there was nonetheless substantial gene expression heterogeneity across the large stromal cell cluster. We find three subtypes of fibroblasts: in addition to the bulk of fibroblasts, we find a myoid fibroblast population expressing *Tpm2*, *Myh11*, and other muscle markers, as well as a distinctive *Angptl7*-positive population of fibroblasts (**Figure S2**). *Angptl7*-positive fibroblasts were previously identified as a subepithelial fibroblast subpopulation in single cell studies of the uterine endometrium, and are predicted to play roles in antigen presentation and regulation of immune responses ^27^. Altogether, we confirm the presence of a fairly typical repertoire of stromal and support cells in the seminal vesicle.

### Secretory epithelial cells

We next turn to the secretory epithelial cells which are primarily responsible for producing the bulk of the fluid secreted by the seminal vesicles. Seminal vesicle secretory epithelial cells are readily identified by high level expression of genes encoding well-known seminal vesicle proteins: *Svs1*, *Svs2*, *Svs5*, and so forth (**Figure 1C**). Perhaps the most surprising marker of this cell population is *Dnase2b*, a relatively understudied class II DNase best known for its role in clearing cell free DNA during lens development ^28^. That said, high level expression of *Dnase2b* has been observed both in a prior RNA-seq survey of the seminal vesicle ^13^, and the encoded protein is abundant in proteomic analyses of murine seminal vesicles and seminal vesicle fluid ^9,12,16,17,19^.

To further explore the potential for functionally-specialized secretory cells in the seminal vesicle, we extracted *Svs*-positive cell populations (**Figure 1A**) for reclustering. Overall we find a relatively homogeneous population (**Figure 2A**), with two notable subclusters (**Figure 2B**). Although the primary population did not separate into distinct clusters, we note some continuous variation across the cells of this main cluster, with cells expressing a gradient of *Svs5* and other SV markers anticorrelated with a gradient of *Atf3* expression (**Figure 2C**). Turning to the two small subclusters, we identified a small population of cells (red cluster) marked by expression of various cell cycle-related genes (*Cenpa*, *Tubb5*, *Top2a*, *Cdk1*, *Mki67*), which presumably represent actively dividing epithelial cells (**Figure 2**). More interestingly, we noted a sizable population of cells (blue cluster) expressing markers (*Tgm4*, *Pbsn*, *Pate9*) typical of the predominant luminal cell population in the prostate ^29^.

**Figure 2.**
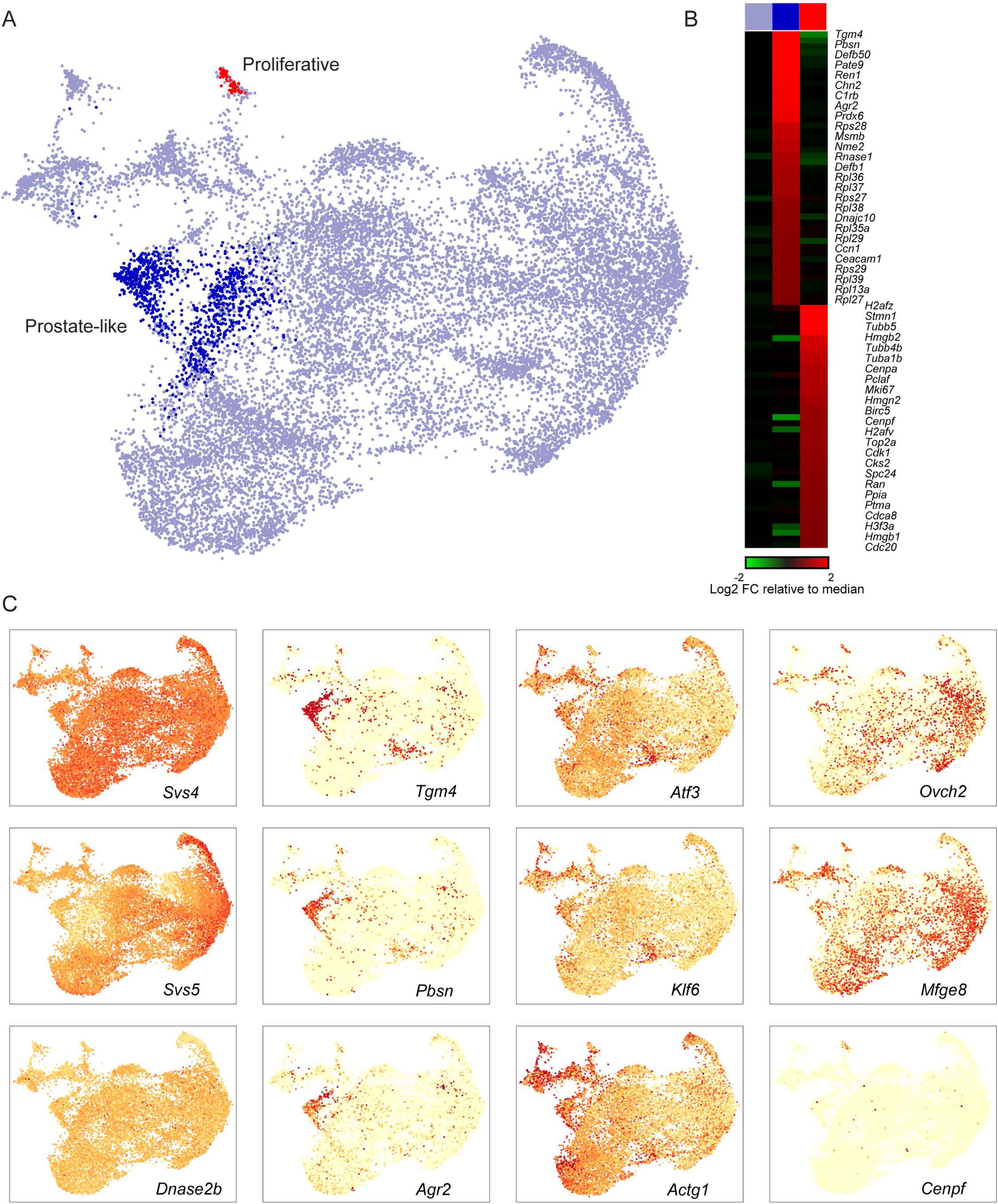
Limited diversity among secretory epithelial cells. A) Reclustering of secretory epithelial cells. Inset shows full dataset, with the cells used for reclustering highlighted in black. Main panel shows a UMAP visualization of reclustered epithelial cells, with two subtypes of epithelial cell labeled. B) Heatmap showing markers for the two minor subclasses of seminal vesicle epithelial cells. C) UMAPs colored according to expression of marker genes for major epithelial cell types.

Given the tight anatomical apposition of the seminal vesicle and prostate, we initially expected that these *Tgm4*+ cells might represent contamination arising during our tissue dissections. However, five considerations led us to speculate that these might be bona fide cell populations in the seminal vesicle itself. First, these cells were consistently observed across the 23 SV samples used in this study, spanning multiple distinct dissections by multiple experimentalists. Second, any low level of contamination by the prostate would primarily arise from the external surface of the prostate, with most cells presumably being stromal rather than epithelial in origin. Third, prior RNA-seq ^13^ and proteomic ^12^ analyses of the seminal vesicles revealed very high levels of the RNA and protein markers of this cell type, with for instance *Tgm4* RNA being the 23^rd^ most abundant transcript in Skerrett-Byrne *et al*, at nearly 10% the levels of the massively-expressed *Svs* genes. This would not be expected from a small amount of contamination arising during dissection. Finally, TGM4, PBSN, and other proteins produced by these cells have been documented in very high levels in pure seminal fluid expressed from the SV ^9,19^, again arguing that these cells are likely to occur in the bona fide SV epithelium. Finally, a Cre driver based on the *Pbsn* promoter was shown to drive recombination not only in the prostate, but also in the seminal vesicle ^30^.

To definitively test the hypothesis that the *Tgm4*+ cells in our seminal vesicle dataset result from prostate contamination, we carried out immunofluorescence for TGM4 in independent seminal vesicle preparations (**Figure S3**). We confirm TGM4 presence both in prostate and in the seminal vesicle, although unexpectedly we find that TGM4 protein staining is not confined to a small subpopulaton of epithelial cells (as expected from the small fraction of *Tgm*4+cells in our RNA-seq dataset), instead finding TGM4 throughout the seminal vesicle epithelium (**Figure S3B**). We therefore visualized *Tgm4* mRNA and pre-mRNA by hybridization chain reaction ^31^ in prostate and seminal vesicle (**Figures S3C-D**). As observed for TGM4 protein, we detected *Tgm4* pre-mRNA and mature mRNA (albeit with relatively weak signal) throughout the seminal vesicle epithelium.

Altogether, our results confirm the expression of *Tgm4* in cells of the seminal vesicle. Beyond this, we cannot find evidence for a distinct subpopulation of *Tgm4*+ epithelial cells in the seminal vesicle as documented in our single cell RNA-Seq dataset (**Figure 2**); our results are instead more consistent with global low level *Tgm4* expression throughout the seminal vesicle (**Figures 3B, D**), along with possible contamination of our single cell dataset by prostate epithelial cells expressing much higher levels of *Tgm4*.

**Figure 3.**
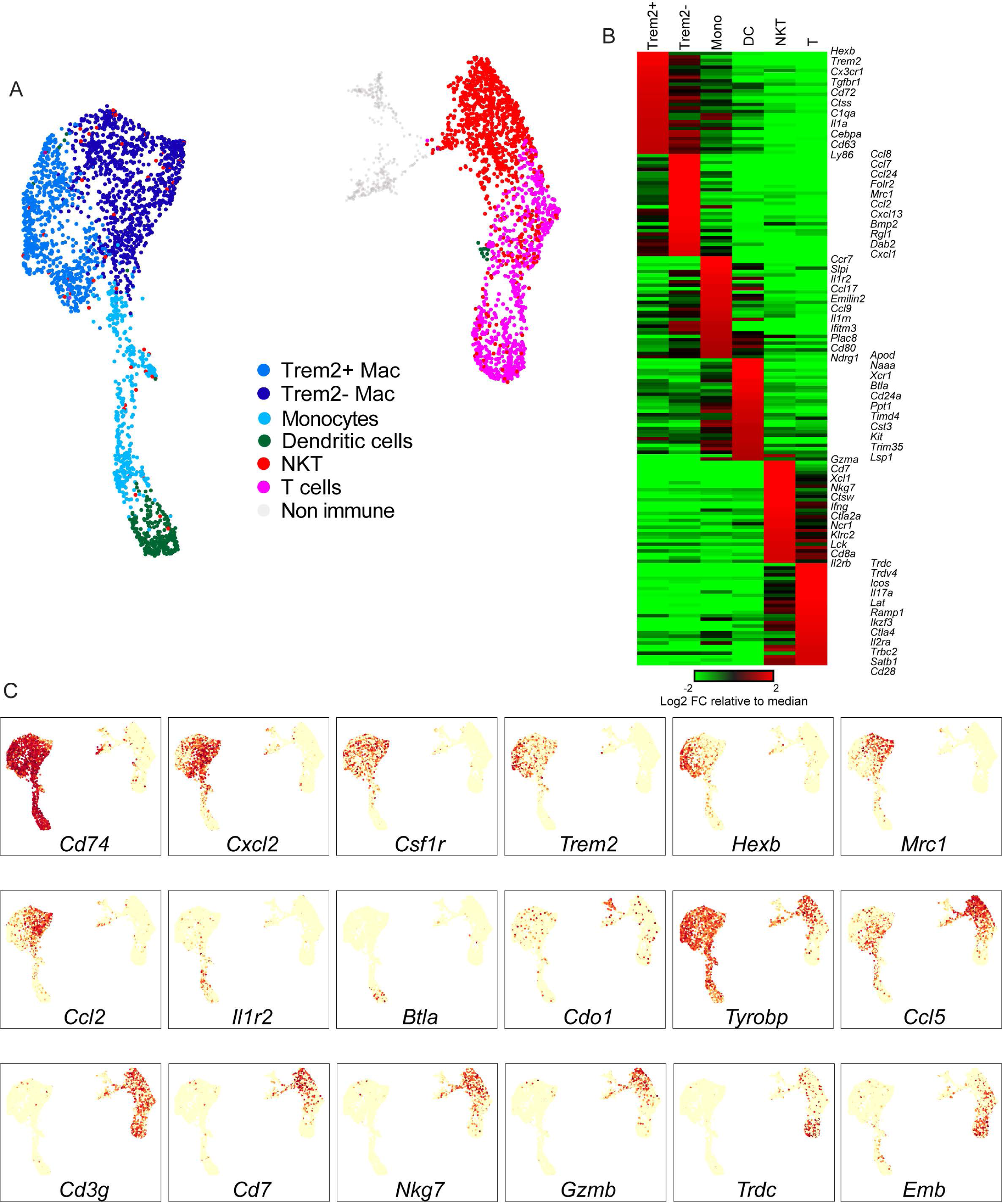
Immune populations of the seminal vesicle. A) Reclustering of seminal vesicle immune populations, as in Figure 2A. B) Heatmap of markers for the indicated immune cell clusters. C) UMAPs colored according to marker genes of interest, as in Figures 1C, 2C.

### Immune populations

Long thought to simply provide a nutritive transport medium for sperm delivery, seminal fluid is now understood to carry a wide array of signaling molecules that prepare the female reproductive tract for fertilization, implantation, and fetal development ^1–5^. In mammals, many of these signaling molecules are cytokines and other immune modulators, raising the question of which cell types secrete these factors into the seminal vesicle fluid – it is typically assumed that many of these molecules are produced by the secretory epithelial cells lining the lumen of the seminal vesicle, but we find that the genes encoding most of these cytokines are undetectable in this cell type (**Figure S4** and **Table S2**). We therefore extracted all presumptive immune populations from the full dataset and reclustered as described above. After reclustering, we recover two broad groups of immune cells – Cd74+MHCII+ antigen presenting cells and CD3+ T/NKT cells – with substantial heterogeneity within each major immune population (**Figure 3A, Table S3**).

Focusing first on the large cluster of *Cd74*/MHC II-positive antigen presenting cells, we identify four distinct populations. Two of these appear to be bona fide macrophages expressing *Csf1r*, *Il1a*, *Adgre1* (aka F4/80), *Cd68*, and *Itgam*, with *Trem2*-positive (*Cd72*, *Hexb*, *Ctsd*, *Tgfbr1*) and -negative (*Mrc1*, *Ccl8*, *Ccl7*, *Cd163*, *Bmp2*) subgroups (**Figure 3B**). Another group of APCs was annotated as dendritic cells based on expression of markers including *Xcr1*, *Wdfy4*, *Itgae* (aka CD103), *Clec9a*, and *Naaa*, among others. Finally, a somewhat nebulous group of MHC II-positive cells expressing *Ccl22* and *Emilin2* – but also sharing markers with macrophages (*Il1b*, *Il1rn*, *Ccl9*) and dendritic cells (*Il1r2*, *Csf2rb*) – were provisionally annotated as monocytes. The second major cluster of immune cells broadly expressed T cell markers including *Cd3d/e/g*, *Il2rb*, and *Trbc1/2*. This group included two substantial subpopulations corresponding to likely NKT cells (*Nkg7*, *Cd7*, *Klrc1/2*) and expressing various cytotoxic effector genes (*Gzma*, *Gzmb*, *Ifng*), along with a population of T cells (*Ctla4*, *Icos*, *Emb*) including both alpha beta (*Trbc1*, *Trbc2*) and delta gamma (*Trdc*, *Trdv4*) T cells.

To further confirm the spectrum of immune cells present in the seminal vesicle, we characterized CD45+ cells by flow cytometry (**Figure 4A**). Consistent with our single cell data, we confirmed the presence of abundant F4/80+ macrophages (40.475%), NKp46+ NK cells (35.55%), and T cells (both CD4+ (8.275%) and CD8+ cells (10.25%)), along with a smaller population of F4/80-CD11c+ dendritic cells (7.075%). Neutrophils (CD11b+ Ly6G+), eosinophils (SiglecF+), and B cells (B220+) could not be detected. Further assessment of the F4/80+ macrophage population demonstrated that the majority of these cells were CD64+CD11c+ (91.8%) (**Figure S5**), with Ly6C+CD11b+ (67.9%) or Ly6C-CD11b+ (16.6%) being the most prominent populations, while a smaller proportion were Ly6C+CD11b-(10.6%) or Ly6C-CD11b-(4.9%) (**Figure 4B**). Histological studies confirmed the presence of both HEXB+ and CD206+ subpopulations of macrophages, both of which were often found closely associated with the secretory epithelium (**Figure 4C**). Together, our data reveal the presence of a complex immune ecosystem in the murine seminal vesicle with the potential to produce a range of cytokines and other immune modulators.

**Figure 4.**
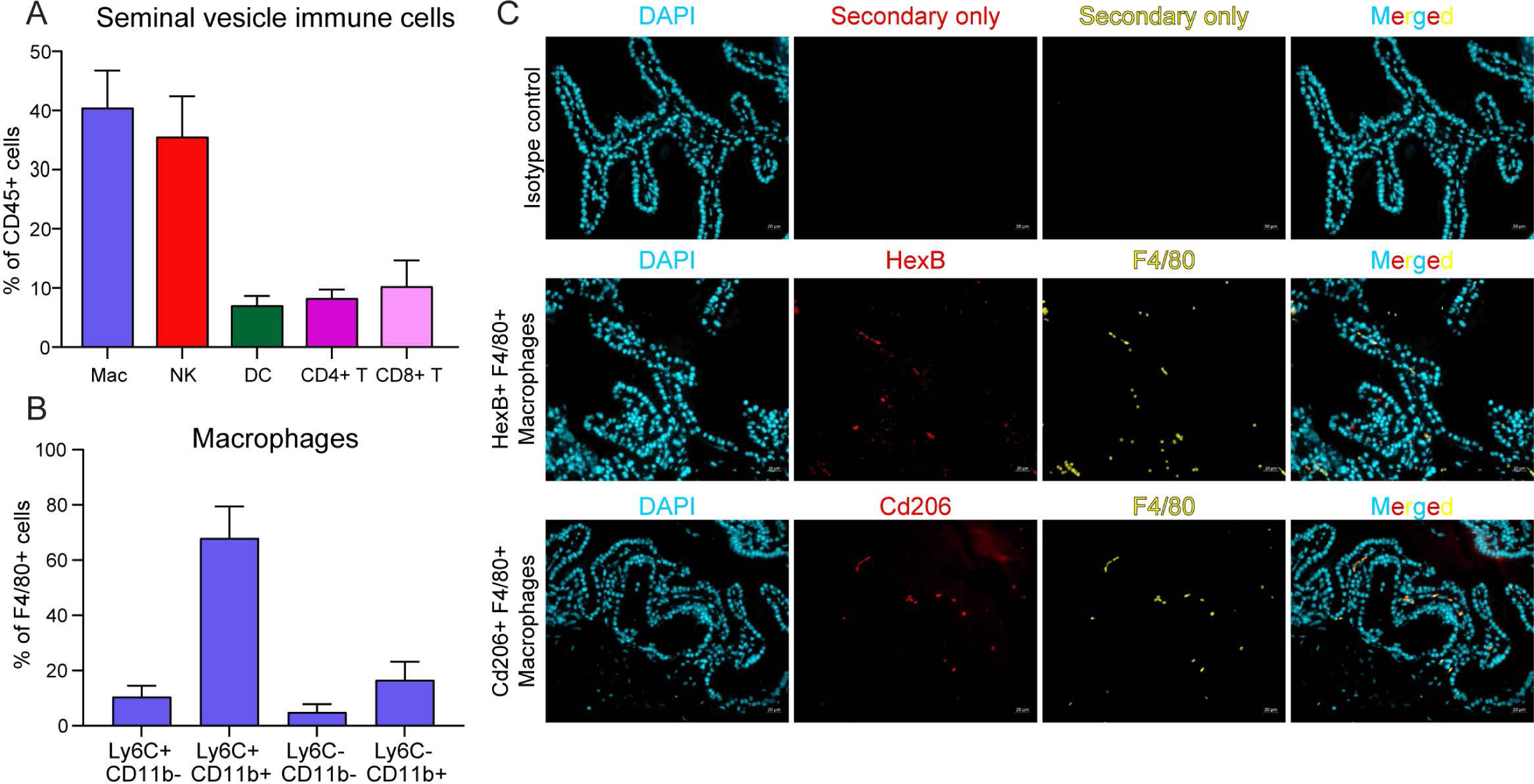
Flow cytometry and immunofluorescence characterization of immune cell populations in the mouse seminal vesicle. Seminal vesicle tissue was collected from 8-12 week old adult male Swiss mice and processed for either (A-B) flow cytometry, or (C) Immunofluorescence. A) Flow cytometry was initially used to confirm single cell RNA-sequencing observations through assessment of Macrophages (Mac), Natural Killer T (NK) cells, Dendritic cells (DCs), and T cells (CD4+ and CD8+). B) Macrophages were further sub-divided into sub-populations through the assessment of LY6C and CD11B. C) Macrophage subtypes identified in scRNA-sequencing were assessed using immunofluorescence. Representative images show F4/80+ macrophages (yellow) in mouse seminal vesicle co-localised with HEXB (middle) or CD206 (bottom) in mouse seminal vesicle tissue. DAPI (cyan) was used as a nuclear counter stain. Images are presented as DAPI alone, HEXB/CD206 alone, F4/80 alone or merged. Isotype controls (top) were used as negative control. Scale bar – 20 mm.

## DISCUSSION

We present here a single cell atlas of the seminal vesicle, one of the least-studied organs in mammals. We recover all the expected cell types in this tissue, and our data provide a resource of RNA expression for all the cell populations. Beyond its utility as a resource, our data reveal two important features of seminal vesicle cell composition.

First, we find relatively little heterogeneity among the primary secretory epithelial cells that produce the bulk of seminal fluid. This stands in contrast to, say, the epididymis ^32–34^, where epithelial cell heterogeneity along the length of this tissue drives the production of molecularly distinct microenvironments experienced by sperm during the days-long maturation process ^35^. In contrast, seminal fluid is secreted into a large single lumen, obviating the need for specialized epithelial cells.

That said, we find three potential sources of seminal vesicle epithelial heterogeneity. First, we identify a small population of epithelial cells expressing a range of cell cycle markers, presumably marking cells undergoing active proliferation. Second, we find a broad continuum of cells across the major secretory epithelial cluster, with cells expressing a gradient of *Svs5* and other major SV markers, anticorrelated with a gradient of *Atf3* expression (**Figure 2C**, compare *Svs5* and *Atf3* profiles). Finally, we identify a subpopulation of epithelial cells expressing *Tgm4*, *Pbsn*, and other well-known markers of the prostate epithelium ^29^. Although multiple lines of evidence suggest that these could be a subpopulation of the epithelial cells of the seminal vesicle – from the presence of TGM4 protein in fluid collected from the SV lumen to the consistent occurrence of these cells across tens of independent dissections – we were unable to identify high-expressing cells in situ by TGM4 protein and RNA staining (**Figure S3**). Instead, we observe modest *Tgm4* RNA and protein expression throughout the seminal vesicle, without any evidence for a subpopulation of high-expressing cells across the many histological sections examined. Thus, although we cannot definitively rule out the presence of such cells in intact SV, our data together are best explained by consistent contamination of SV dissections with cells from the tightly-adhered prostate tissue.

The second key observation in this study is the absence of cytokine expression in the secretory epithelial cells that produce the bulk of seminal fluid. Cytokines and other immune modulators are emerging as key components of seminal fluid which help to shape the maternal immune response to sperm and to the developing embryo, and thus represent key signaling molecules for mammalian reproduction. The absence of most cytokines in the RNA profile of the epithelial cells thus suggests the hypothesis that seminal fluid cytokines might be produced by infiltrating immune cells in the seminal vesicle. This hypothesis will be tested in future studies, as understanding the cellular origin of seminal fluid signaling molecules is essential for defining pathways by which diet and stress challenges impact male to female signaling.

Altogether, our data provide a valuable resource for the reproductive biology community. This dataset will serve as a baseline for future studies focused on the effects of androgen signaling, dietary challenges, and other stressors on seminal vesicle gene regulation and secretory function.

## ACKNOWLEDGEMENTS

We thank B. Nixon, S. Robertson, and members of the Rando lab for insightful discussions and comments on the manuscript. This work was funded by NIH R01AG073238 (OJR), University of Newcastle College of Engineering, Science and Environment External Collaboration, and Fellowship Accelerator grants (JS), and NIA grants (VNG).

## MATERIALS AND METHODS

### Animal husbandry

Male FVB/NJ and C57BL/6J mice at age of about 12 weeks to 28 months were used in this study. The animals were maintained under controlled temperature and humidity conditions on a 12 hr light/dark cycle with different diets and water ad libitum. The diets used included Chow diet (product # 5P76), or three diets based on AIN-93G: Control diet (product # 3156, CD), High Fat diet (product # 3282, HFD) or Low Protein diet (product # 4579, LP). The mice on HFD, LP, and CD were fed for 9 weeks after weaning until sacrifice at 12 weeks of age. For caloric restriction, singly-housed animals were provided with 70% of the mass of AIN-93G consumed by a singly-house animal consuming diet ad libitum. All animal care and use procedures were in accordance with guidelines of the University of Massachusetts Medical School Institutional Animal Care and Use Committee (Protocol # 201200029).

For flow cytometry and immunofluorescence experiments, male outbred Swiss mice (8–12 weeks of age) were obtained from a breeding colony held at the University of Newcastle central animal facility and maintained according to the recommendations prescribed by the Animal Care and Ethics Committee. Mice were housed under a controlled lighting regimen (12-h light: 12 h dark) at 21 to 22 °C and supplied with food and water ad libitum.

### Seminal vesicle dissection and single cell dissociation

Seminal vesicles were collected from 12-week to 28-month old male mice (**Table S1**). In brief, after euthanizing the mice, the urogenital system was exposed. The entire urogenital system was taken out of the mouse abdominal cavity and placed into a 60 mm Petri dish containing 5 ml complete medium (CM) composed of IMDM (Invitrogen), 10% FBS (Sigma), and Antibotic-Antimycotic (invitrogen). Surrounding organs and tissues were carefully removed under a dissection microscope, and the seminal vesicle was opened longitudinally with a scissor to let seminal vesicle fluid out of the organ. The tissue was then transferred to another dish and any loose coagulated seminal vesicle fluid was removed without disturbing the epithelial cell layer. After washing, the tissue was transferred into a 15 ml conical tube with 2 ml CM and 2 ml extracellular matrix digestion media (ECM) containing collagenase (Sigma), DNase (Sigma), collagenase/hyaluronidase (Stem Cell), and collagenase/dispase (Sigma). The tissue was incubated in a water bath at 300 rpm for 30 minutes at 35 C. After incubation, 10 ml IMDM-DNase media was added to the tube and centrifuged at 300 g for 5 minutes at room temperature. The pellet was washed once with 8 ml IMDM-DNase media, resuspended with 3 ml TrypLEtm express (Gibco), and digested in the water bath at 300 rpm for 5 minutes at 35 C. The tube was then filled with 10 ml CM and the cell suspension was filtered through a series of 100 μm, 70 μm, and 40 μm cell strainers.

Cells were pelleted by centrifuging for 5 minutes at 300 g, and the pellet was resuspended with 3 ml CM. The cell suspension was then placed on the top of OptitPrep (Sigma) gradient followed by centrifuging at 800 g for 15 minutes. The gradient was composed of 3 ml of 67%, 50%, and 36% OptiPrep in CM, respectively. Interface fractions were collected and cells were pelleted, washed 1X with CM, and resuspended in 1 ml CM. Only samples with cell viability over 80% were used for sequencing library preparation.

### Single cell RNA-seq

Single-cell sequencing libraries were prepared using Chromium Single Cell 3’ Reagent Kit V2 (10X Genomics), as per the manual, and sequenced on NextSeq 500 and HiSeq 4000 at the UMass Medical School Deep Sequencing Core.

### Data analysis

Seurat was used for clustering single-cell data. We filtered cells that have a number of unique features (genes) less than 500, a number of unique molecular identifiers (UMI) less than 200, and a percentage of mitochondrial genes over 1%. After filtering, we have total 47,804 cells and 19,601 features (genes) with average 6053 UMI per cell. Filtered data was normalized with SCTransform as described in ^36^.

Transformed data across conditions were then integrated with canonical correlation analysis. Genes from the FindVariableFeatures function were used as input for the initial principal component analysis (PCA). The number of principal components (PCs) was chosen based on the Seurat ElbowPlot function. We set resolution parameters from 0.1 to 1.4 and examined the clustering quality based on DoHeatmap function and clustree. We identified significantly differentially expressed markers in each cluster by running the default FindAllMarkers function. We annotated cell types based on well-established markers. After initial annotation, we extracted all major cell types for reclustering in **Figures 2**, **3**, **S2**.

Data will be submitted to GEO during the review process.

### Flow cytometry staining and analysis

The procedure for obtaining single cell suspensions was loosely based on previous published protocols for obtaining single cell uterine tissue ^37^, and further refined empirically. Seminal vesicle tissue from male mice was dissected ensuring that the anterior prostate (coagulating gland) and other fact and connective tissue was left in situ. Seminal vesicle fluid was allowed to flow from the gland prior to storage of the tissue in a petri dish containing ice-cold HEPES buffer (10mM HEPES-NaOH pH 7.4, 150mM NaCl, 5mM KCl, 1mM MgCl2, 1.8mM CaCl2) and placed on ice. Tissue was submerged in HEPES buffer, cut into smaller pieces using sharp pointed scissors prior to the addition of 2 mg/mL Collagenase D (ThermoFisher Scientific) and 80U/mL DNase I (Roche). Tissue fragments were then digested with gentley shaking at 37°C for 30 minutes using a gentleMACS Tissue Dissociator (Miltenyi Biotec, Macquarie Park, Australia). Following dissociation, cells were filtered through a 70 μM cell strainer followed by a wash in HEPES buffer. Cells were then pelleted at 500 x g for 10 minutes at 4°C before being incubated in red blood cell lysis buffer (155 mM NH4Cl, 12mM NaHCO_3_, 0.1mM ethylenediaminetetraacetic acid [EDTA], pH 7.35) for 5 minutes at 4°C. Red blood cell lysis was stopped by the addition of an equal amount of FACS buffer (2% fetal bovine serum [FBS], 2mM EDTA in PBS) before cells were centrifuged at 500 x g for 10 minutes at 4°C. Cells were then resuspended in FACS buffer and counted prior to staining for flow cytometry.

Cells were then incubated in 10ng/mL Fc bBlock (anti-mouse CD16/CD32; BioXCell) for 15 minutes. Panels of directly conjugated antibodies as indicated in Supplemental Table X were subsequently added and incubated on ice for a further 30 min. Cells were washed twice with FACS buffer and resuspended in FACS buffer for immediate assessment on a Fortessa X20 flow cytometer (BD Biosciences). The results were analysed using FACS Diva software (BD Biosciences), gating only on viable cells and excluding red blood cells. Gating strategy is shown in **Figure S5**.

### Immunofluorescence staining and analysis

For immunofluorescence analysis of macrophages (**Figure 4C**), excised seminal vesicles from the gland containing seminal vesicle fluid were fixed for 8 h in Bouin’s solution (5% v/v acetic acid, 25% v/v formaldehyde, 70% v/v picric acid). Bouin’s solution was removed with daily changes of 75% (v/v) ethanol for at least 5 days.

Tissues were embedded in paraffin and 5 μm sections were cut prior to staining. Sections were dewaxed in xylene, followed by rehydration in ethanol and water. Slides were subjected to antigen retrieval in a pressure cooker using a 10 mM sodium citrate, pH 6 solution (for 10 minutes at ∼60Kpa and 110^0^C -118^0^C). After a rinse in MilliQ H2O, endogenous peroxidases were blocked with 3% hydrogen peroxide for 30 minutes, followed by a TBST (50 mM Tris-HCl, 150 mM NaCl, 0.1% Tween-20 in distilled H2O, pH 7.6) wash (3 x 5 min). Subsequently, sections were incubated in a solution of 5% BSA and 10% goat serum in TBST for 30 minutes at room temperature to prevent nonspecific antibody binding. All antibodies were subsequently diluted in 2.5% BSA and 5% goat serum in TBST. The slides were then incubated with primary antibodies HEXB (diluted 1:100, PA5-101082, Thermofisher Scientific) or CD206/MRC1 (diluted 1:50, MA5-16871, Thermofisher Scientific), along with appropriate isotype controls, overnight at 4°C. After incubation, slides were washed in TBST (3 x 5 min) and incubated with either anti-rabbit or anti-rat secondary antibodies with AF594 (diluted 1:400, A11012 or A1107, Thermofisher Scientific) for 1 hour. Following another TBST wash (3 x 5 min) and a second antigen retrieval step at 37°C using proteinase K (20 μg/ml in TBS, pH 8, V3021, Promega), the slides were blocked as described above. This was followed by a second primary antibody incubation: F4/80 (diluted 1:50, 14-4801-82, Thermofisher Scientific) overnight at 4°C. Following TBST washes (3 x 5 min), the slides were incubated with an anti-rat secondary antibody AF488 (diluted 1:400, A11006, Thermofisher scientific) for 1 hour, followed by TBST washes (3 x 5 min) and then counterstaining with DAPI (20µg/ml in TBST, D9542, Sigma Aldrich). Slides were then mounted using an antifade reagent composed of 10% v/v Mowiol 4 to 88, 30% v/v glycerol in 0.2 M Tris (pH 8.5), and 2.5% v/v 1,4-diazobicyclo-(2.2.2)-octane in 0.2 M Tris (pH 8.5). Imaging and photography were conducted using a Zeiss Axio A.2 fluorescence microscope (Carl Zeiss AG).

For TGM4 immunostaining in prostate and seminal vesicle (**Figures S3A-B**), seminal vesicles were collected and fixed with 4% paraformaldehyde (PFA)/PBS or Bouin’s at 4C overnight. After washing the excess of PFA/Bouin’s with PBS, sectioning was done at 5 μm thickness by the UMASS morphology core. Slides were stained as recommended by the antibody manufacturer. Briefly, slides were progressively dewaxed using Xylene washes, then rehydrated through a series of ethanol solutions.

Permeabilization with 0.1% Tween-20 in PBS was performed prior to a 1 hr blocking with a blocking solution containing 10% goat serum (Vector Laboratories) and 3% BSA (Sigma). The primary and secondary antibodies specifications and dilutions are at 1:500 and 1:2000, respectively. All slides were incubated with primary antibodies at 4°C overnight, washed 3 X 5 min with PBST and subsequently incubated with suitable Alexa Fluor secondary antibodies for 1 hr at room temperature, followed by a 3 x 5 min washes with PBST. Slides were mounted with VECTASHIELD PLUS Antifade Mounting Medium with DAPI (Vector Laboratories) and imaged the following day using a Zeiss Axio inverted microscope. Images were then processed to increase brightness and contrast using Photoshop.

### Hybridization chain reaction.

PFA fixed seminal vesicle were prepared and sectioned as above. DNA probes sets and DNA HCR amplifiers were purchased from Molecular Instruments. The sequences used for probe design were transcript ENSMUST00000026893.6 for mature *Tgm4* transcript and the intron 1-2 for pre-*Tgm4* transcript. The experiment was performed following manufacturer’s manual. Briefly, after a series of dewaxinization, the tissue sections were rehydrated through a serious of ethanol solutions. The sections were pretreated with citrate buffer (pH at 6.0) in microwave for 15 min. After cooling to room temperature, the slides were incubated in PBST at room temperature for 10 min. Prehybridization was carried out in a humidified box at 37C for 10 min, followed by hybridization at 37 C overnight. After three washes with probe wash buffer and 5 X SSCT, signal amplification was performed with hybridization chain reaction (HCR) at room temperature in a humidified box overnight. After four washes with 5 X SSCT, Slides were mounted with VECTASHIELD PLUS Antifade Mounting Medium with DAPI (Vector Laboratories) and imaged with a Zeiss Axio inverted microscope and Nikon confocal microscope. Images were then processed to increase brightness and contrast using Fiji software.

## SUPPLEMENTARY FIGURES AND TABLES

**Figures S1-S5 Tables S1-S3**

## SUPPLEMENTARY FIGURE LEGENDS

**Figure S1.**
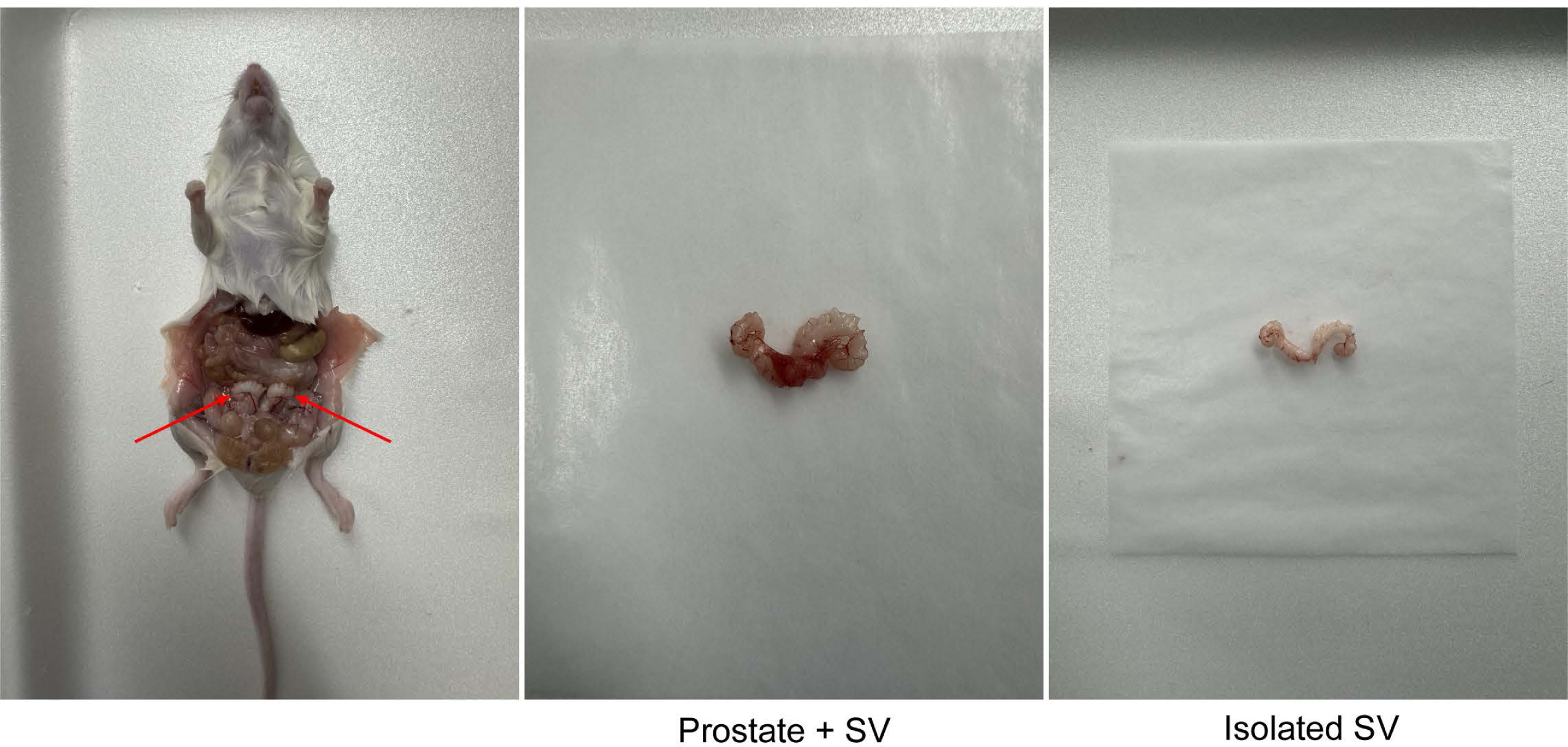
Seminal vesicle dissection. Seminal vesicle from a typical dissection for single cell dissociation. Red arrows on left panel show seminal vesicle in situ, followed by excised seminal vesicle and prostate, and then cleaned seminal vesicle in the rightmost panel.

**Figure S2.**
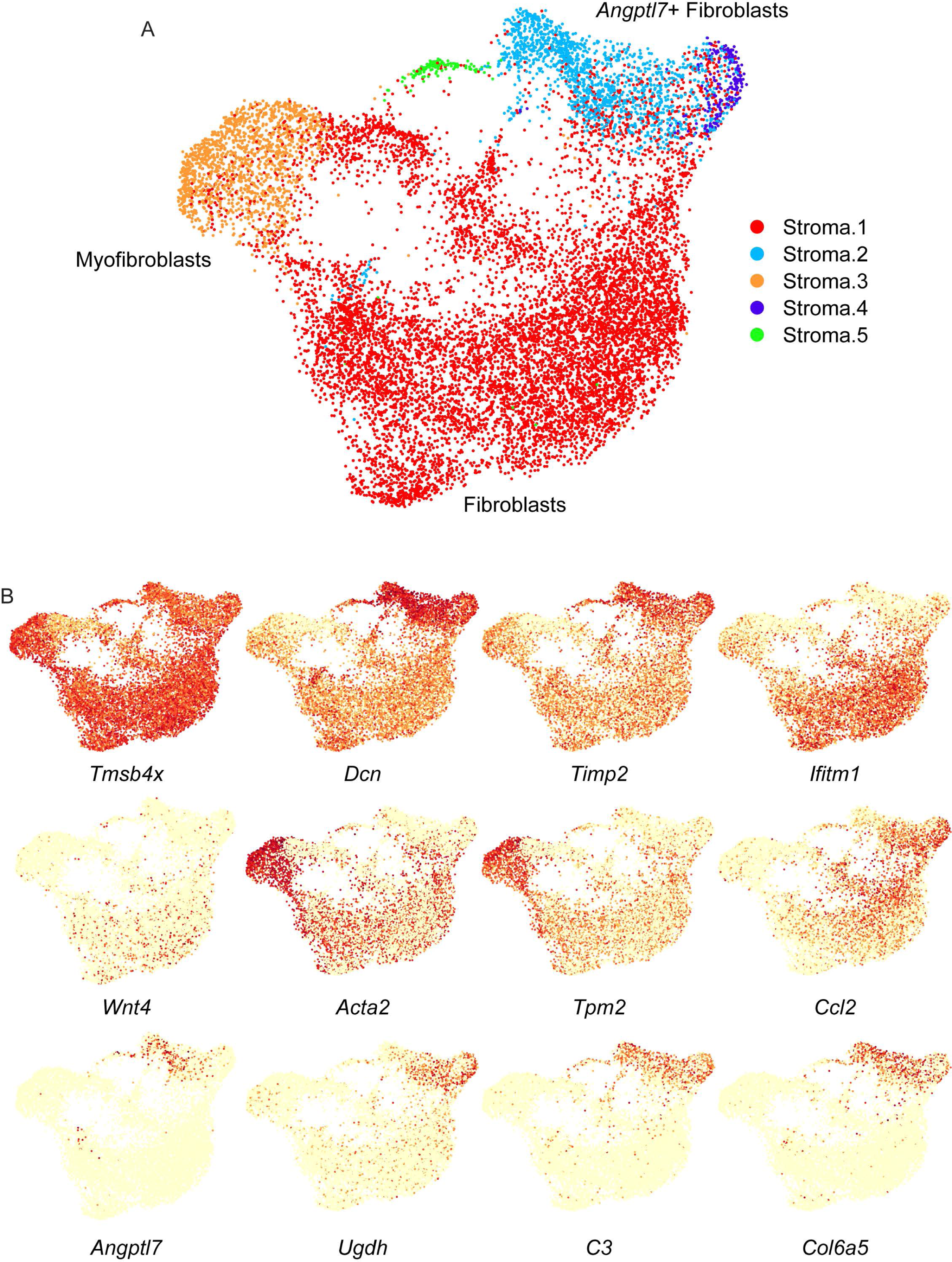
Stromal cell reclustering. A) UMAP shows reclustered stromal cell populations. Inset shows the full dataset, highlighting the clusters extracted for reclustering. Reclustered cells are annotated according to inferred cell type. B) Expression of the indicated genes in the reclustered stromal cell UMAP.

**Figure S3.**
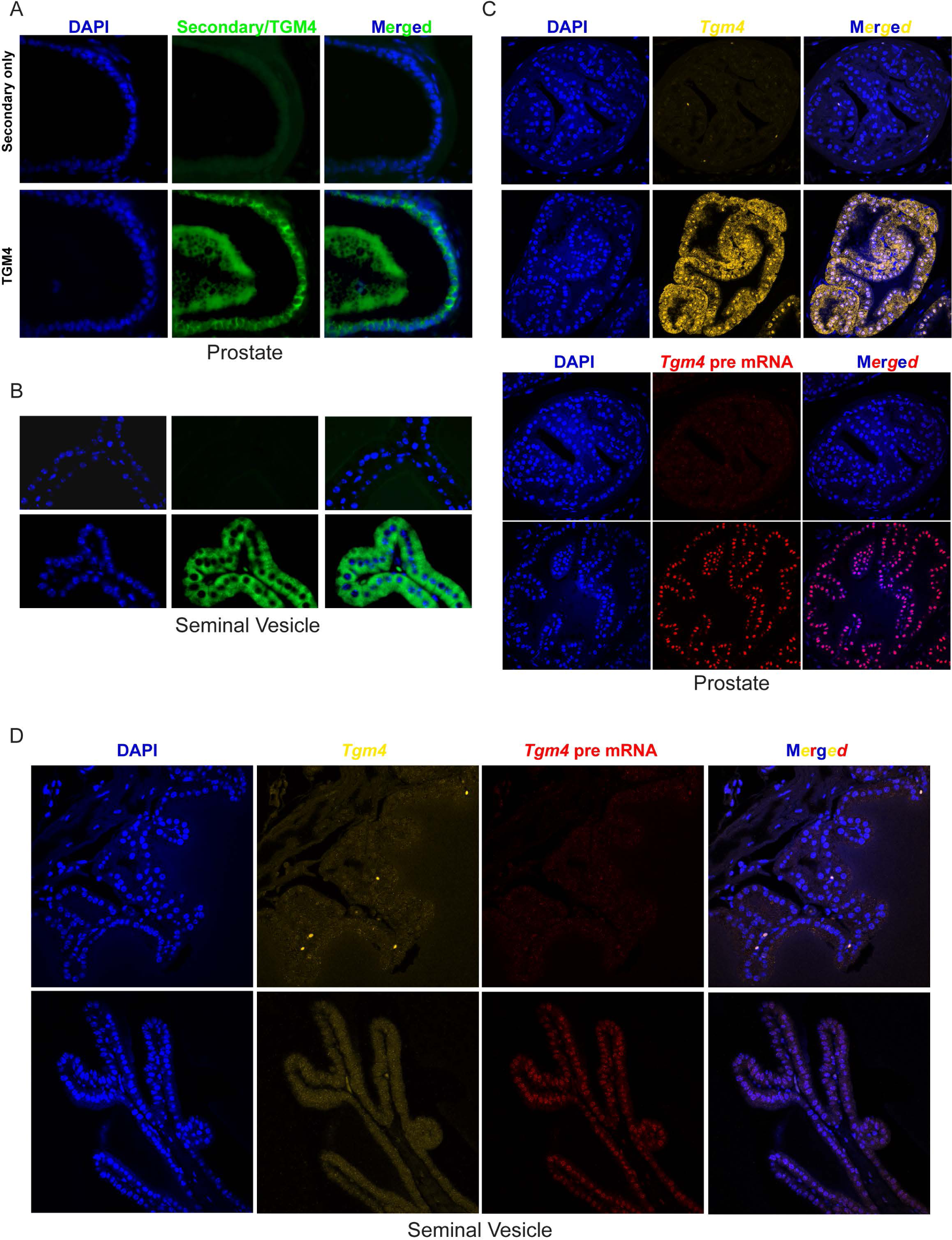
Tgm4 protein and RNA expression in the seminal vesicle. A-B) TGM4 protein expression in the prostate (A) and seminal vesicle (B). Top panels shows secondary antibody only, bottom shows prostate section stained with anti-TGM4. C-D) *Tgm4* RNA expression in prostate (C) and seminal vesicle (D). Hybridization chain reaction probes against spliced *Tgm4* mRNA, or against intron-containing pre-mRNA, as indicated.

**Figure S4.**
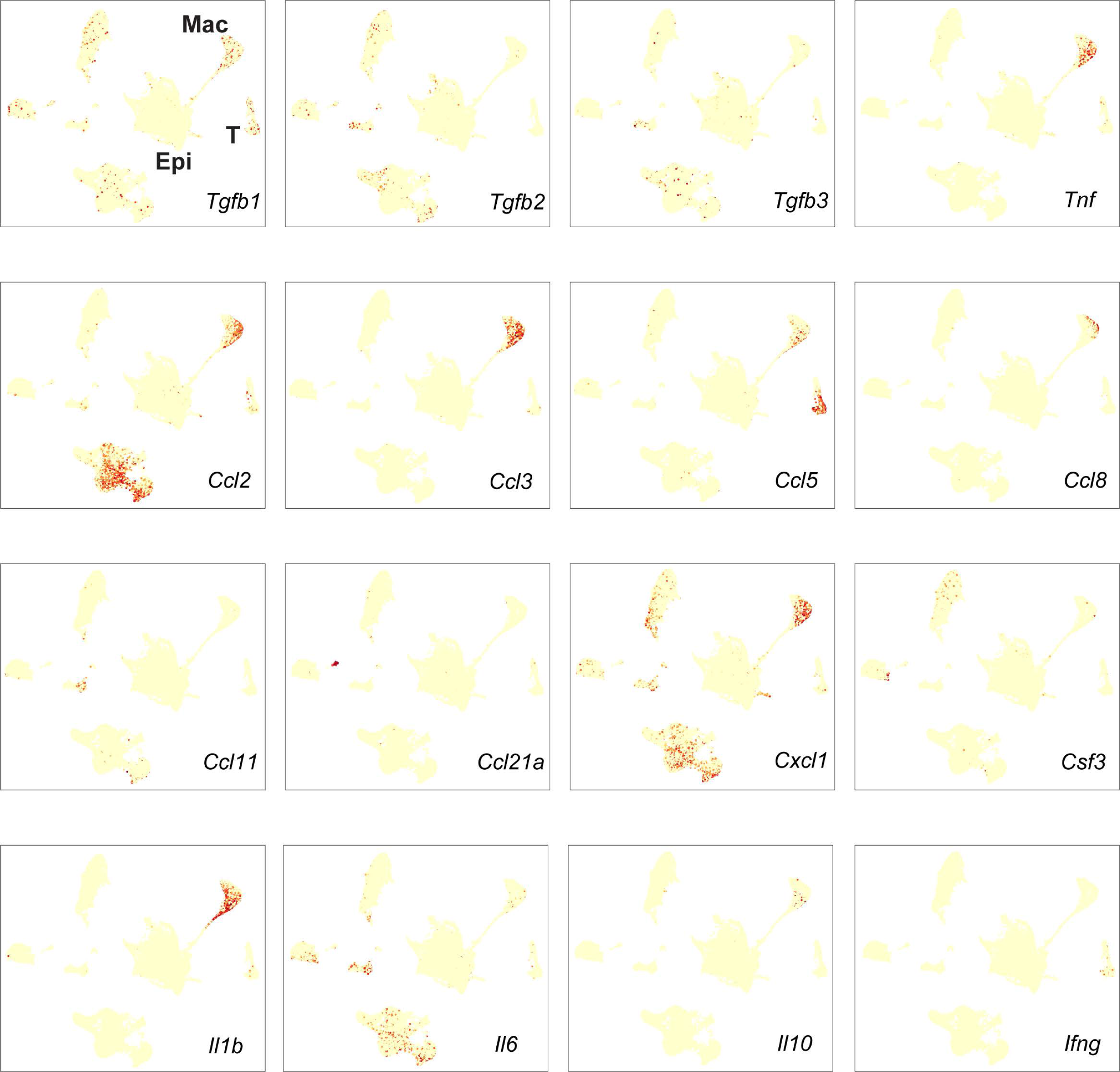
Most cytokines detected in seminal fluid are not detectably expressed in seminal vesicle epithelial cells. UMAPs showing expression of various cytokines previously detected in seminal plasma and linked to seminal fluid signaling in the female reproductive tract. Upper left panel highlights secretory epithelial cells (Epi), macrophages (Mac), and T/NK cells (T) from Figure 1A.

**Figure S5.**
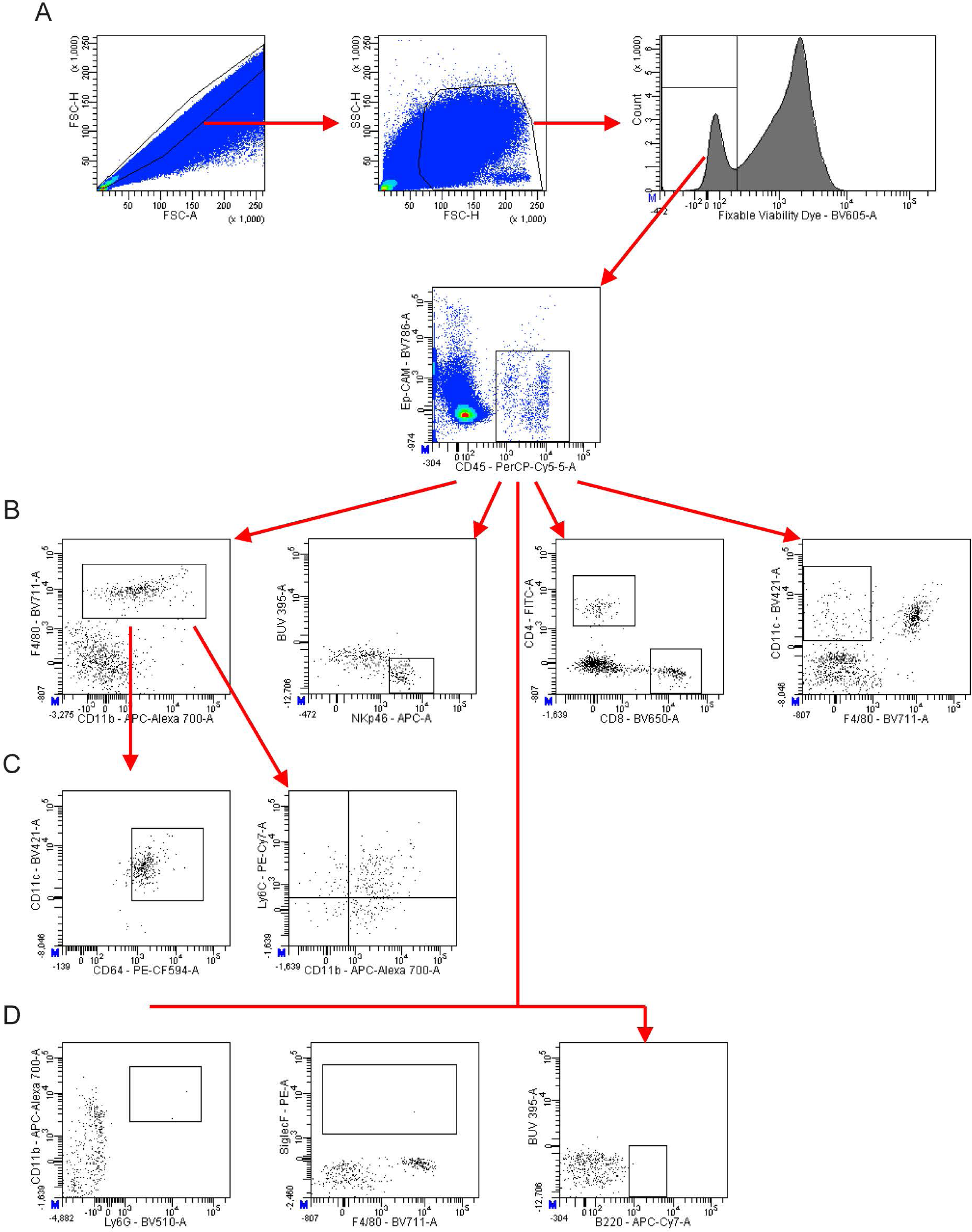
Flow cytometry gating strategy and representative immune cell populations identified in the murine seminal vesicles. Seminal vesicle tissue was collected from 8-12 week old adult male Swiss mice and processed for flow cytometry identifying immune cells residing in seminal vesicle tissue. A) Singlets and intact total cells were selected based on forward scatter (FSC) and side scatter (SSC) followed by live cells (viability dye negative) and leukocytes (CD45+ Ep-CAM-cells). B) Gating and representative frequencies of macrophages (F480+), natural killer (NK) cells (NKp46+), CD4^+^ T cells (CD4+), CD8^+^ T cells (CD8+), and dendritic cells (DCs; CD11c+ F4/80-). C) To determine the phenotypes of macrophages present, F4/80+ cells were gated for CD64, CD11c, CD11b and Ly6C expression. D) Gating for neutrophils (CD11b+ Ly6G+), eosinophils (SiglecF+), and B cells (B220+). Data information: A-D: representative flow cytometry dot plots from one experiment n=4 (biological replicates). B, C: % displayed are the proportions of the parent populations the cells within the gates/quadrants comprise.

## SUPPLEMENTARY TABLES

**Table S1. Animals used for single cell dataset**

Demographic information, including age and dietary exposures, for the animals used for the seminal vesicle samples throughout this study.

**Table S2. Gene expression for cell populations in the murine seminal vesicle**

RNA abundance for each of the major seminal vesicle cell clusters shown in Figure 1A.

**Table S3. Gene expression for immune cell subclustering**

RNA abundance for each of the immune cell clusters shown in Figure 3A.

## Notes

### Competing Interest Statement

The authors have declared no competing interest.

